# Evidence for flexible motor costs in vertical arm movements: reduced gravity-related effort minimization under high accuracy constraints

**DOI:** 10.1101/2024.03.27.586943

**Authors:** Gabriel Poirier, Damien De Filippis, Cyril Sirandre, Charalambos Papaxanthis, Jeremie Gaveau

## Abstract

The central nervous system (CNS) is thought to use motor strategies that minimize several criteria, such as end-point variability or effort, to plan optimal motor patterns. In the case of vertical arm movements, a large body of literature demonstrated that the brain uses a motor strategy that takes advantage of the mechanical effects of gravity to minimize muscle effort. Results from other studies suggested that the relative importance of each criterion may vary according to the task’s constraints. For example, it could be hypothesized that reduced end-point variability driven by high accuracy demands is detrimental to effort minimization. The present study probes this specific hypothesis using the framework of gravity-related effort minimization. We asked twenty young healthy participants to perform vertical arm reaching movements towards targets whose size varied across conditions. We recorded the arm kinematics and electromyographic activities of the anterior deltoid to study two well-known motor signatures of the gravity-related optimization process; i.e., directional asymmetries on velocity profiles and negative epochs on phasic muscular activities. The results showed that both indices were reduced as target size decreased, demonstrating that the gravity-related optimization process was reduced under high accuracy constraints. This phenomenon is consistent with the use of a trade-off strategy between effort and end-point variability. More generally, it suggests that the CNS is able to appropriately modulate the relative importance of varied motor costs when facing varying task demands.

## Introduction

Flexibility is a major feature of the human motor behavior, allowing individuals to plan appropriate movements in varying environments. It is, therefore, essential to understand how motor strategies adapt to varied tasks requirements and characteristics of the environment (Liu and Todorov, 2007). A prominent approach to explain motor strategies is the optimal control theory (Wolpert, 1997; Jordan and Wolpert, 1999; Scott, 2004; Todorov, 2004; Franklin and Wolpert, 2011). Within this framework, the motor system plans movements that minimize varied motor costs – i.e., effort, end-point variability or jerk, for instance – according to the task at hand (Jordan and Wolpert, 1999).

However, the lack of evidence in favor of a single unifying optimization criteria led researchers to the idea that the central nervous system (CNS) might use a weighting of several criteria (i.e., composite cost) to plan optimal motor patterns (Liu and Todorov, 2007; Gielen, 2009; Berret et al., 2011). Recent investigations using inverse optimal control approaches supported the existence of such composite costs (Liu and Todorov, 2007; Mombaur et al., 2010; Berret et al., 2011; Berret and Jean, 2016; Vu et al., 2016). Notably, Berret et al. (2011) used direct and inverse optimal control to determine the optimal cost (among several kinematic, dynamic, and effort costs) that best fit participants’ motor behavior during an arm reaching task with target redundancy. The results revealed that a composite cost mixing kinematic (joint smoothness) and effort (mechanical or neural effort) explained the observed kinematics more accurately than any other single cost. Other works confirmed that such models, minimizing a composite effort/kinematic cost, could appropriately capture human motor patterns during arm point-to-point movements (Gaveau et al., 2014, 2016, 2021), motor learning in a velocity-dependent force field (Healy et al., 2023), and arm raising while standing upright (Tanis et al., 2023).

The concept of composite costs can help understand human motor flexibility. According to this concept, the relative importance of each cost would be modulated according to task requirements. In this spirit, the study of Liu and Todorov (2007) supports a modulation of feedback gains according to task constraints. However, to our knowledge, no study has provided evidence of a modulation of the weighting of optimization costs according to task demand. A straightforward prediction is that effort optimization should be reduced in a highly constrained task compared to an ecological one where the CNS could exploit redundant task dimensions to save muscle effort; i.e. movement duration or end-point position.

Gravity-related motor planning (see Gaveau et al., 2016, 2021) could represent a useful paradigm to address this question. Multiple studies investigating vertical arm reaching movements consistently reported direction-dependent arm kinematics: the time to peak acceleration and time to peak velocity were shorter for upward than for downward movements (Papaxanthis et al., 2005; Gentili et al., 2007; Le Seac’h and McIntyre, 2007; Gaveau and Papaxanthis, 2011; Gaveau et al., 2014; Yamamoto and Kushiro, 2014; Gaveau et al., 2016, 2021; Yamamoto et al., 2016, 2019; Hondzinski et al., 2016; Poirier et al., 2023b, 2023a, 2020, 2022). Optimal control simulations have explained these directional differences - and their progressive disappearance during microgravity exposure (Gaveau et al., 2016) - as the signature of a gravity-related optimization process that minimizes muscle effort. More precisely, these features were predicted by a model minimizing a composite cost combining effort (the absolute work of torques) and jerk (Smooth-Effort model; Berret et al., 2008; Gaveau et al., 2014, 2016, 2021). Another noteworthy prediction of this model was the existence of specific negative epochs on the phasic activity of antigravity muscles during vertical arm movements (Gaveau et al., 2021). Because the tonic electromyogram (EMG)—i.e., the muscle activity that is hypothesized to compensate for gravity torque—is subtracted from the full EMG signal to obtain phasic EMG (Flanders and Herrmann, 1992; Buneo et al., 1994; Flanders et al., 1994), negative phasic EMGs mean that muscular activity is not sufficient to compensate gravity torque and thus that gravity torque participates in the arm’s motion (Gaveau et al., 2021). The existence of these negative phases has been experimentally observed in the deceleration phase of upward movements and in the acceleration phase of downward movements, thereby supporting the model’s predictions (Gaveau et al., 2021; Poirier et al., 2022, 2023b, 2023a). Consequently, directional asymmetries and negatives epochs on phasic EMG are thought to represent the hallmark of the gravity-related effort-optimization process.

Some results suggest that the gravity-related effort optimization process may be influenced by task’s demands (speed instructions notably; (Gaveau and Papaxanthis, 2011; Poirier et al., 2023a). However, no study has hitherto specifically investigated this hypothesis. In arm reaching tasks, the accuracy requirement has long been shown to strongly affect motor strategies (Fitts, 1954; Soechting, 1984; Gribble et al., 2003). A well-documented strategy in the presence of high accuracy constraints is the effortful increased agonist-antagonist coactivation, leading to increased joint stiffness and thus reduced end-point variability (Gribble et al., 2003; Osu et al., 2004; Missenard et al., 2008; Missenard and Fernandez, 2011). Within the optimal control framework, this result can be interpreted as a reduction of the effort cost weighting in the context of a high accuracy requirement.

Here, building upon multiple results supporting a gravity-related effort optimization process, we probed the hypothesis that the effort cost weighting changes with varying accuracy constraints. To this aim, we compared gravity-related effort optimization signatures (i.e., directional asymmetries and negative phases) between vertical reaching movements to varying target sizes. We expected the size of these markers to be reduced for smaller target sizes, that would demonstrate a downweighing of the effort cost (Gaveau et al., 2016) in the presence of high accuracy demands.

## Materials and Methods

### Participants

Twenty healthy young adults [15 males and 5 women; mean age = 25 ± 3 (SD) years] participated in the present experiment after giving their written informed consent. All participants had normal or corrected-to-normal vision without any neurological or muscular disorders. All participants were right-handed, as attested by a laterality index superior to 60 (Edinburgh Handedness Inventory, Oldfield 1971). A French National ethics committee (2019-A01558-49) approved all the experimental procedures. The research was conducted following legal requirements and international norms (Declaration of Helsinki, 1964).

### Experimental protocol

The experimental protocol was similar to previous studies investigating gravity-related arm movement control (Gentili et al., 2007; Gaveau and Papaxanthis, 2011a; Gaveau et al., 2014, 2016; Poirier et al., 2020; Gaveau et al., 2021; Poirier et al., 2022, 2023b). Specifically, participants were asked to perform unilateral single degree-of-freedom vertical arm pointing movements (rotation around the shoulder joint) to a visual target in a parasagittal plane with their dominant arm (Fig. 1A). During these movements, the work of gravity torque varies with movement direction, whereas inertia remains constant. Thus, such movements are known to provide an appropriate experimental paradigm to investigate gravity-related motor planning. To maximally vary the effect of gravity torque on the movement, participants moved in two opposed directions: upward and downward.

**Figure. 1.**
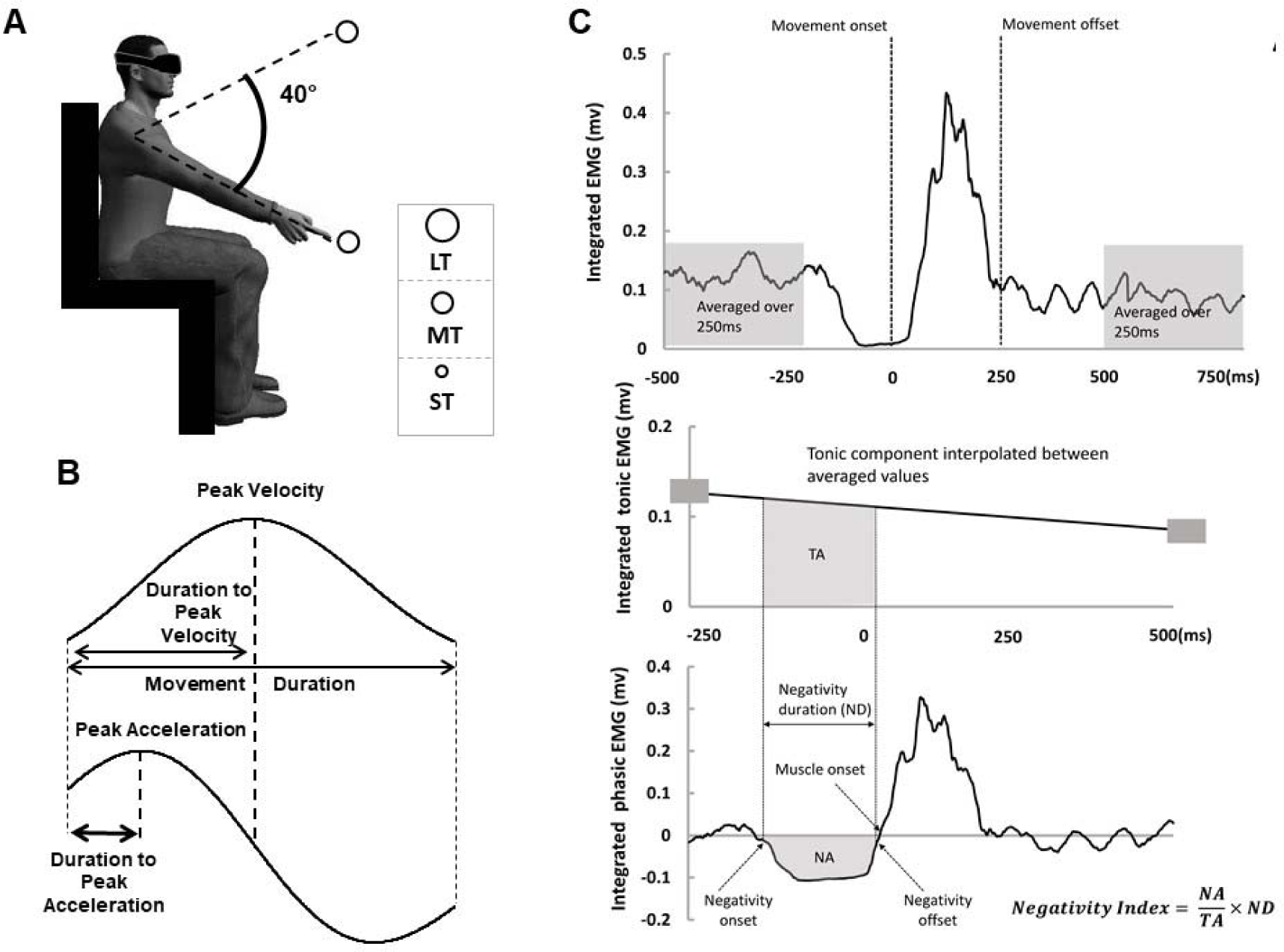
A. Experimental setup. Participants performed arm pointing movements between two virtual targets whose size varied across conditions. B. Illustration of the parameters computed on the velocity and acceleration profiles. C. Analysis of the tonic and phasic components of the EMG. Illustration of the integrated full EMG signal. Movement onset and offset were detected on the velocity profile. Upper panel: We averaged the signal from 500 to 250ms before movement onset and from 250 to 500ms after movement offset (grey windows). Middle panel: Tonic EMG component was obtained by linear interpolation between the two previously averaged values (small grey rectangles). Lower panel: Phasic EMG component was obtained by subtracting the tonic component (in middle) from the full EMG (in upper panel). We detected negativity onset and offset and then computed negativity duration (NDur) and integrated phasic signal between negativity onset and offset (NA). TA in panel B represents the integrated tonic signal in the same period.

We examined the effect of accuracy constraint on gravity-related motor planning by varying target size among blocks. For this purpose, participants were immersed in a virtual reality environment through a head-mounted display (HTC VIVE; Fig. 1A). Our virtual reality environment was developed using Unity software and consisted of a basic outdoor urban environment that provided visual gravity clues (e.g. buildings) to the participant. The targets (colored spheres) were displayed in this environment. Participants were provided real-time feedbacks concerning their hand’s position in the environment during entire pointing movement.

The experiment was conducted in a block design. We asked participants to execute vertical pointing movements in three conditions (blocks): to a large (LT; target diameter = 70mm), a medium (MT; diameter = 50mm) and a small (ST; diameter = 30mm) target (Fig.1A). Participants carried out 32 movements whose directions were randomly interleaved within each block. Overall, each participant performed 96 trials (16 trials x 2 directions x 3 target sizes) throughout the experiment.

Participants sat on a chair with their trunk parallel to the gravity vector. Two targets were presented in the parasagittal plane crossing the participant’s right shoulder. Targets were displayed at a distance corresponding to the length of the participant’s fully extended arm plus two centimeters. Vertical positions of targets were tuned for each participant to require a 40° shoulder flexion or extension (Fig. 1A). Targets were centered around the antero-posterior horizontal line crossing the shoulder joint, thereby corresponding to a 110° (upward target) and 70° (downward target) shoulder elevation.

A trial was carried out as follows: the participant positioned her/his fully extended arm in front of the initial target (verbally indicated by the experimenter using a color code). After a brief delay (∼2 seconds), the experimenter informed the participant that she/he was free to reach to the other target, whenever she/he wanted. The participant had been informed that the reaction time and the movement velocity were not constrained. At the end of a movement, participants were requested to maintain their final position (about 2 seconds) until the experimenter instructed them that they were free to relax their arm. Participants were invited to have a rest between trials (∼10 s) and blocks (5mn) to prevent muscle fatigue, and potential ocular fatigue and dizziness due to the virtual reality environment. Before each block, participants were invited to perform few practice trials (∼5 trials) to familiarize with the task.

We placed five reflective markers on the participant’s shoulder (acromion), arm (middle of the humeral bone), elbow (lateral epicondyle), wrist (cubitus styloid process), and finger (nail of the index). We also recorded spatial location of both targets. Three-dimensional position was recorded using an optoelectronic motion capture system (Vicon system, Oxford, UK; six cameras; 200Hz). After calibration, spatial variable error of the system was less than 0.5mm. Additionally, we recorded muscle activation patterns by means of bipolar surface electrodes positioned over the anterior (AD) and posterior (AD) heads of deltoid muscle (Aurion, Zerowire EMG, sampling frequency: 1000Hz). The Giganet unit (Vicon, Oxford, UK) allowed synchronized recording of kinematic and EMG signals.

### Data analysis

We processed Kinematic and EMG data using custom programs written in Matlab (Mathworks, Natick, NA). Data processing was similar to previous studies (Poirier et al., 2020, 2022; 2023; Gaveau et al., 2021).

#### Kinematics

Markers’ position was filtered (third-order low-pass Butterworth filter; 5Hz cut-off, zero-phase distortion, “butter” and “filtfilt” functions) before computing velocity and acceleration profiles by numerical differentiation (3 points derivative). We computed angular joint displacements to ensure that elbow and wrist rotations were negligible (<1° for each trial). Movement onset and offset were respectively defined as the times when finger velocity rose above and fall below a 10% of peak velocity threshold. Movements presenting more than one local maxima, i.e., more than one velocity peak, were automatically rejected from further analysis (less than 2%). The following parameters were computed to quantify kinematic patterns (Fig. 1B): 1) Movement duration (MD = offset - onset). 2) Relative duration to peak acceleration (rD-PA = duration to peak acceleration divided by MD). 3) Relative duration to peak velocity (rD-PV = duration to peak velocity divided by MD). 4) Distance between finger end-point position and target position (spatial error). Because the task was to perform vertical single degree-of-freedom movements, this distance was calculated on the vertical axis only. For each participant and condition, constant and variable errors were calculated as follows:

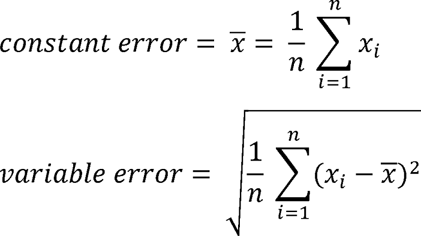

Where n is the total number of trials in the condition (16 for each condition), *x_i_* is the spatial error between finger end-point position and target position for the i^th^ movement of the condition, and *x̅* is the constant error in the condition. We then computed from these variables an index of total variability, representing how successful participants were in achieving the target (Schmidt and Lee, 2019). Total error was computed for each participant and condition according to the formula:

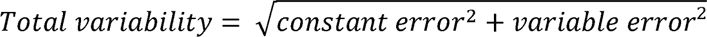

#### Electromyography

We filtered (bandpass third-order Butterworth filter; 20-300Hz, zero-phase distortion, “butter” and “filtfilt” functions) and then rectified EMG signals before integrating them using a 50ms sliding window from 250ms before movement onset to 250ms after movement offset. EMG signal were then normalized by the EMG recorded during a maximal voluntary isometric contraction (MIVC, same EMG processing as movement patterns). At the beginning of each block, participants performed three MIVC in both flexion and extension directions, at a 90° shoulder angle. Lastly, we ordered EMG traces according to movement mean velocity and averaged them across two trials (from the two slowest to the two fastest movements), resulting in 8 EMG traces to be analyzed for each block and direction. Before averaging two trials together, we normalized the duration of each trial to the mean duration value of these two trials.

To emphasize the role of gravity in the arm motion, we separated the phasic and tonic components of each EMG signal using a well-known subtraction procedure (Flanders and Herrmann, 1992; Buneo et al., 1994; Flanders et al., 1994; D’Avella et al., 2006, 2008; Gaveau et al., 2021; Poirier et al., 2022). We computed averaged values of the integrated phasic EMG signals from 500ms to 250ms before movement onset and from 250 to 500ms after movement offset (Fig. 1C). Then, we extracted the theoretical tonic component using a linear interpolation between these two averaged values (Fig. 1C). Finally, we computed the phasic component by subtracting the tonic component from the full EMG signal (Fig. 1C).

It was recently shown that phasic EMG activity of antigravity muscles consistently exhibits negative epochs during vertical arm movements (Gaveau et al., 2021; Poirier et al., 2022; 2023a; 2023b) when the arm’s acceleration sign is coherent to gravity’s – i.e., in the acceleration phase of downward movements and in the deceleration phase of upward movements). Model simulations demonstrated that the negativity of antigravity muscles reflects an optimal motor strategy where gravity force is harvested to save muscle force. We defined negative epochs as a time interval when phasic EMG was inferior to zero minus a 95% confidence interval (computed on the integrated EMG signals from 500ms to 250ms before movement onset) for at least 50ms. We used this value as a threshold to define negativity onset (when phasic EMG dropped below) and offset (when phasic EMG rose above). Then, we computed relative negativity duration (negativity duration divided by movement duration), negativity occurrence – i.e., the number of trials including a negative period divided by the total number of trials in the condition – and a negativity index to compare the amount of negativity between conditions:

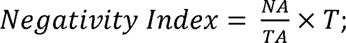

where NA stands for Negative Area and is the integrated phasic signal between negativity onset and offset; TA stands for Tonic Area and is the integrated tonic signal between negativity onset and offset (if NA and TA are equal, it means that the muscle is completely silent during the interval); and T is the duration of the negative epoch normalized by movement duration. Figure. 1C provides an illustration of the computation of EMG parameters.

In addition to those negativity-related parameters, we computed a coactivation index, quantifying the amount of coactivation in each movement, from the agonist and antagonist EMG traces (Latash, 2018). In each trial, the coactivation was calculated at each point in time with the formula: 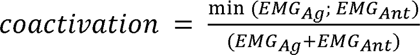, where *EMG_Ag_* and *EMG_Ant_* refer to the values of respectively agonist and antagonist EMG signals. The coactivation index was defined as the mean of these coactivation values over MT. With this index, a value of 0.5 corresponds to maximal coactivation while the value of zero corresponds to no coactivation.

### Statistics

Statistical analyses were performed using JASP. Distribution normality (Kolgomorov-Smirnov test) and sphericity (Mauchly’s test) was checked for all variables. To analyze movement duration and amplitude, we used repeated measure ANOVA with two within participant factor: *target size* (LT vs. MT vs. ST) and *direction* (up vs. down). We used *HSD-Tukey tests* for post-hoc comparisons. We analyzed correlations between variables with repeated measures correlations (rmcorr package in R, Bakdash and Marusich, 2017). The level of significance was set at p<0.05 for all analyses.

## Results

A qualitative illustration of velocity and phasic EMG profiles in all conditions for a typical participant is provided in Fig.2. Table.1 displays the means (±SD) values of all kinematic parameters, for each condition. All participants’ motions lied in the parasagittal plane (shoulder azimuth angle <1° for all trials; *n* = 1730) with bell-shaped, single-peaked velocity profiles (Fig. 2). Generally, all participants were able to accurately reach the target (variable error was inferior than 25mm for all participants in all conditions; absolute constant error was inferior than 15, 25, and 55mm in the ST, MT and LT conditions, respectively, for all participants). In this section, we first describe the reorganization of the movements’ kinematics following target size reduction with previously-used measures (i.e., movement duration, variable error, and constant error). We then investigate the effect of target size on gravity-related motor optimization through directional asymmetries on the velocity profile (i.e., rD-PA and rD-PV).

**Figure.2.**
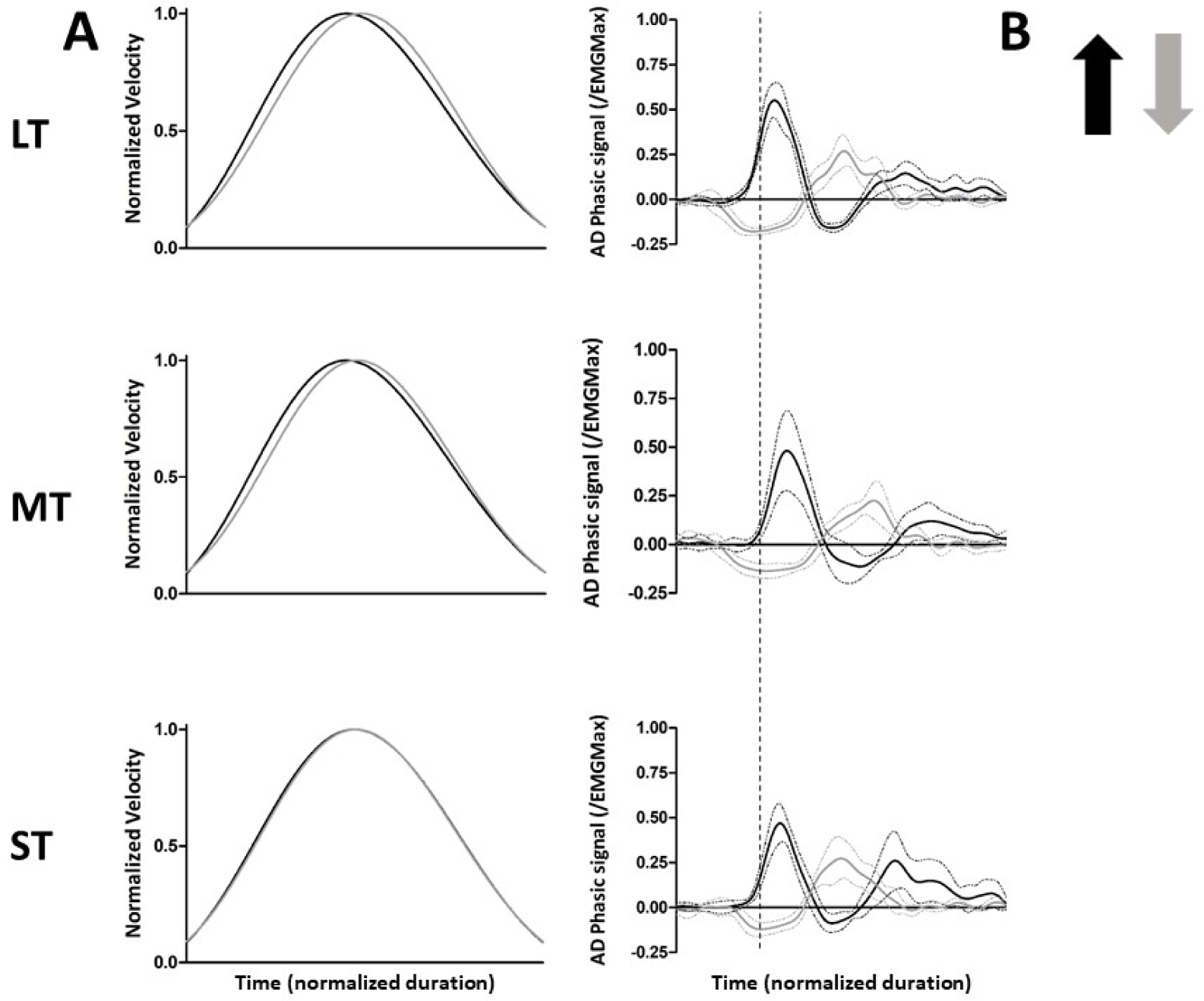
A. Mean velocity profiles of upward (black traces) and downward (grey traces) movements in a typical participant across conditions. B. Illustration of the mean (± SD) phasic EMG patterns in a typical participant across conditions. Dashed lines represent the kinematic onset of the movement. Vertical arrows indicate movement directions.

**Table.1.**
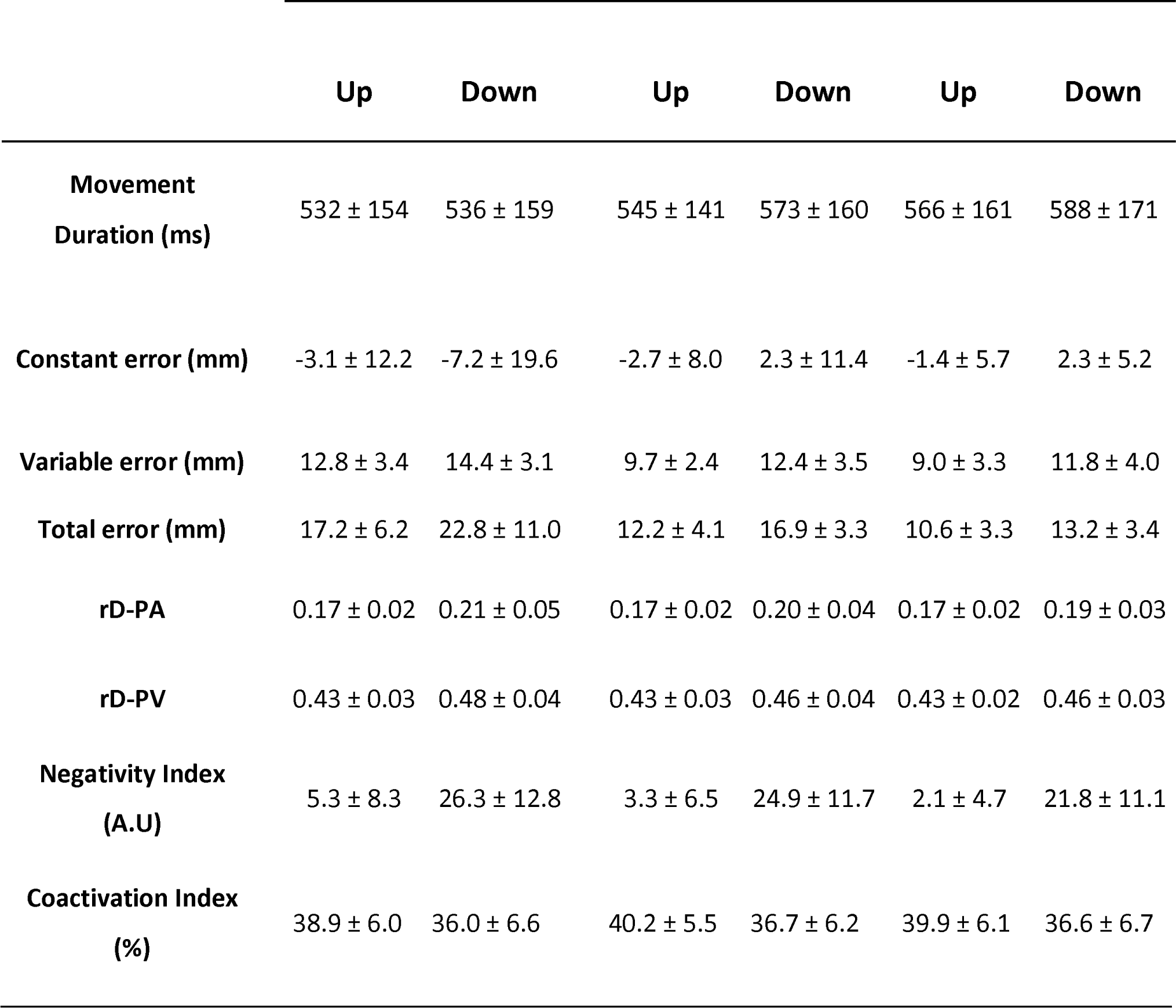
Mean (±SD) values of all kinematic and EMG parameters.

|  | Up | Down | Up | Down | Up | Down |
| --- | --- | --- | --- | --- | --- | --- |
| **Movement**<br>**Duration (ms)** | 532 ± 154 | 536 ± 159 | 545 ± 141 | 573 ± 160 | 566 ± 161 | 588 ± 171 |
| **Constant error (mm)** | -3.1 ± 12.2 | -7.2 ± 19.6 | -2.7 ± 8.0 | 2.3 ± 11.4 | -1.4 ± 5.7 | 2.3 ± 5.2 |
| **Variable error (mm)** | 12.8 ± 3.4 | 14.4 ± 3.1 | 9.7 ± 2.4 | 12.4 ± 3.5 | 9.0 ± 3.3 | 11.8 ± 4.0 |
| **Total error (mm)** | 17.2 ± 6.2 | 22.8 ± 11.0 | 12.2 ± 4.1 | 16.9 ± 3.3 | 10.6 ± 3.3 | 13.2 ± 3.4 |
| **rD-PA** | 0.17 ± 0.02 | 0.21 ± 0.05 | 0.17 ± 0.02 | 0.20 ± 0.04 | 0.17 ± 0.02 | 0.19 ± 0.03 |
| **rD-PV** | 0.43 ± 0.03 | 0.48 ± 0.04 | 0.43 ± 0.03 | 0.46 ± 0.04 | 0.43 ± 0.02 | 0.46 ± 0.03 |
| **Negativity Index**<br>**(A.U)** | 5.3 ± 8.3 | 26.3 ± 12.8 | 3.3 ± 6.5 | 24.9 ± 11.7 | 2.1 ± 4.7 | 21.8 ± 11.1 |
| **Coactivation Index**<br>**(%)** | 38.9 ± 6.0 | 36.0 ± 6.6 | 40.2 ± 5.5 | 36.7 ± 6.2 | 39.9 ± 6.1 | 36.6 ± 6.7 |

### General parameters

#### Movement Duration

Fig. 3A displays mean (± SD) movement duration (MD) for each condition. It is well-known that the duration of pointing movements is influenced by the target’s diameter (Fitts, 1954). Accordingly, we found a significant *target size* effect (F = 5.44; p = 8.36e-3; η_p_^2^ = 0.22), with increased MD as target size decreased (Fig. 3A). Additionally, MD was significantly influenced by *direction* (F = 7.17; p = 1.49e-2; η_p_^2^ = 0.27): participants exhibited longer MD for downward compared to upward movements (Fig. 3A). This difference was significant for ST (*post-hoc*: p = 7.02e-3; Cohen’s d = 0.14) and MT (p = 4.83e-4; Cohen’s d = 0.18), but not in the LT condition (p = 1.70e-1; Cohen’s d = 0.09). *Target size x direction* interaction effect did not reach significance (F = 1.38; p = 2.63e-1; η_p_^2^ = 0.07).

**Figure.3.**
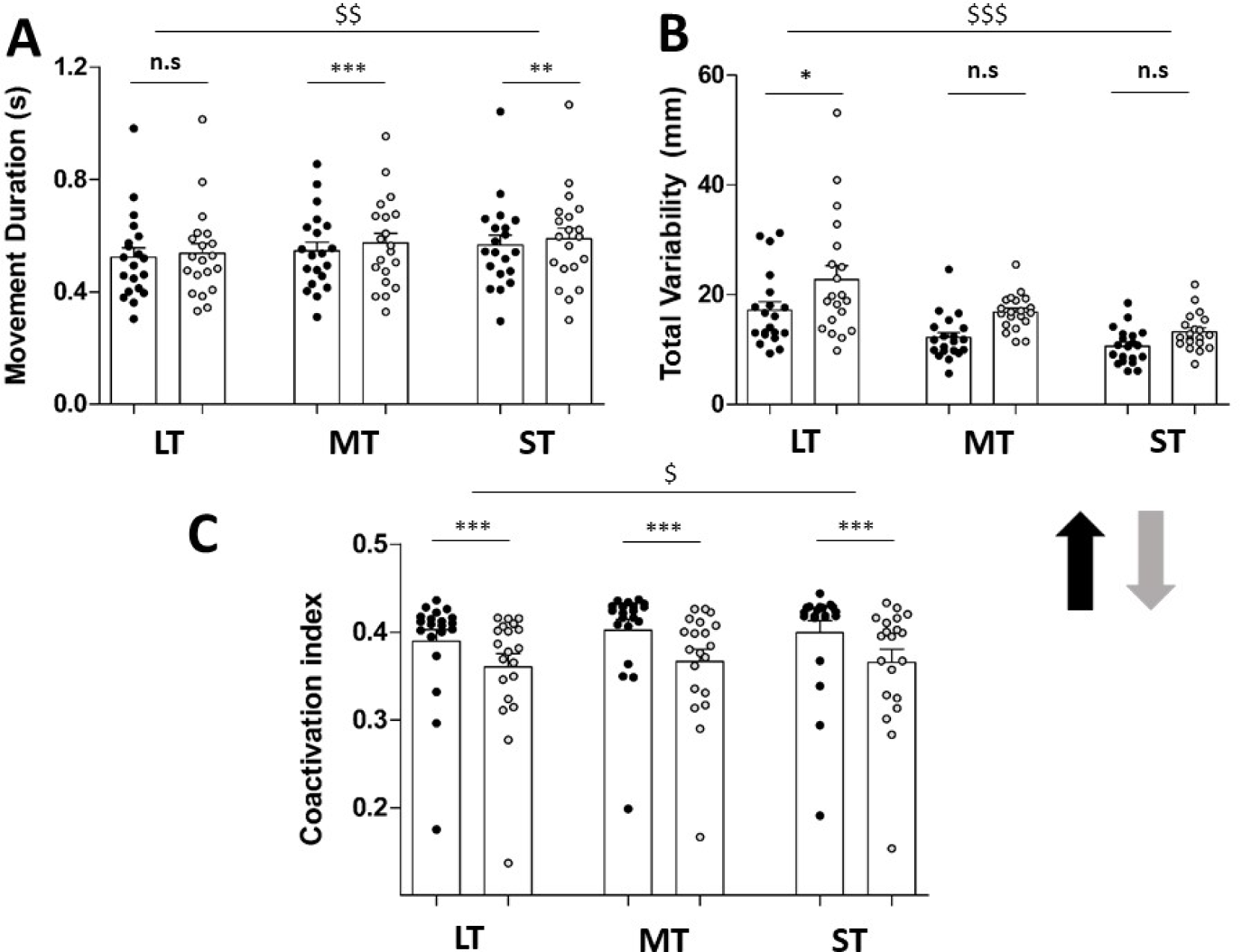
A. Illustration of the mean (±SD) movement duration in upward (black) and downward (grey) movements for all target size conditions. Dots represent individual values of all participants. B. Illustration of the mean (±SD) total variability in upward (black) and downward (grey) movements for all target size conditions. Dots represent individual values of all participants. C. Illustration of the mean (±SD) coactivation index in upward (black) and downward (grey) movements for all target size conditions. Dots represent individual values of all participants. $: Significant main effect of target size; p < 0.05 $$: Significant main effect of target size; p < 0.01 $$$: Significant main effect of target size; p < 0.001 *: upward vs. downward difference on post-hoc test: p < 0.05 **: upward vs. downward difference on post-hoc test: p < 0.01 ***: upward vs. downward difference on post-hoc test: p < 0.001

#### Total Variability

Fig. 3B displays mean (± SD) total variability for each condition. As imposed by task demands, total variability significantly decreased with target size (*target size* effect: F = 19.90; p = 1.22e-6; η_p_^2^ = 0.51; see Fig. 3B). We also observed a significant *direction* effect (F = 14.42; p = 1.21e-3; η_p_^2^ = 0.43). It was highlighted by *post-hoc* comparisons that total variability was significantly larger for downward compared to upward movements in the LT (p = 1.18e-2; Cohen’s d = 0.94) but not in MT (p = 5.44e-2; Cohen’s d = 0.78) and ST (p = 5.71e-1; Cohen’s d = 0.43; see Fig. 3B) conditions. *Target size x direction* interaction effect was not found to be significant (F = 0.97; p = 3.89e-1; η_p_^2^ = 0.05).

#### Coactivation Index

Fig. 3C displays mean (± SD) coactivation index for each condition. Both *target size* (F = 4.37; p = 1.95e-2; η_p_^2^ = 0.19) and *direction* (F = 13.88; p = 1.44e-3; η_p_^2^ = 0.42) effects were found to be significant. As previously observed (Gribble et al., 2003), muscular coactivation was shown to increase for smaller target sizes (Fig. 3C). Additionally, *post-hoc* comparisons revealed that coactivation index was smaller in downward compared to upward movements for all conditions (up vs. down: p < 1.40e-4; Cohen’s d > 0.47 in each case; Fig. 3C). *Target size x direction* interaction effect was not significant (F = 1.36; p = 2.68e-1; η_p_^2^ = 0.07).

### Gravity-related effort optimization parameters

#### Kinematics

##### rD-PA

Mean (± SD) rD-PA in each condition is presented in Fig. 4A. Both *target size* (F = 7.51; p = 1.79e-3; η_p_^2^ = 0.28) and *direction* (F = 50.33; p = 9.51e-7; η_p_^2^ = 0.73) effects were found to significantly influence rD-PA. As previously demonstrated, rD-PA was shorter for upward compared to downward movements in all target size conditions (Fig. 4A; upward vs downward comparisons: p < 6.72e-4; Cohen’s d > 0.66 in each case). In downward movements, rD-PA was longer in the LT condition compared to the ST (p = 2.89e-4; Cohen’s d = 0.72) and the MT (p = 3.15e-2; Cohen’s d = 0.46) conditions but did not statistically differ between the ST and the MT conditions (p = 4.51e-1; Cohen’s d = 0.26). However, there was no difference between target size conditions in upward movements (p > 7.62e-1; Cohen’s d < 0.20 for all comparisons).

**Figure.4.**
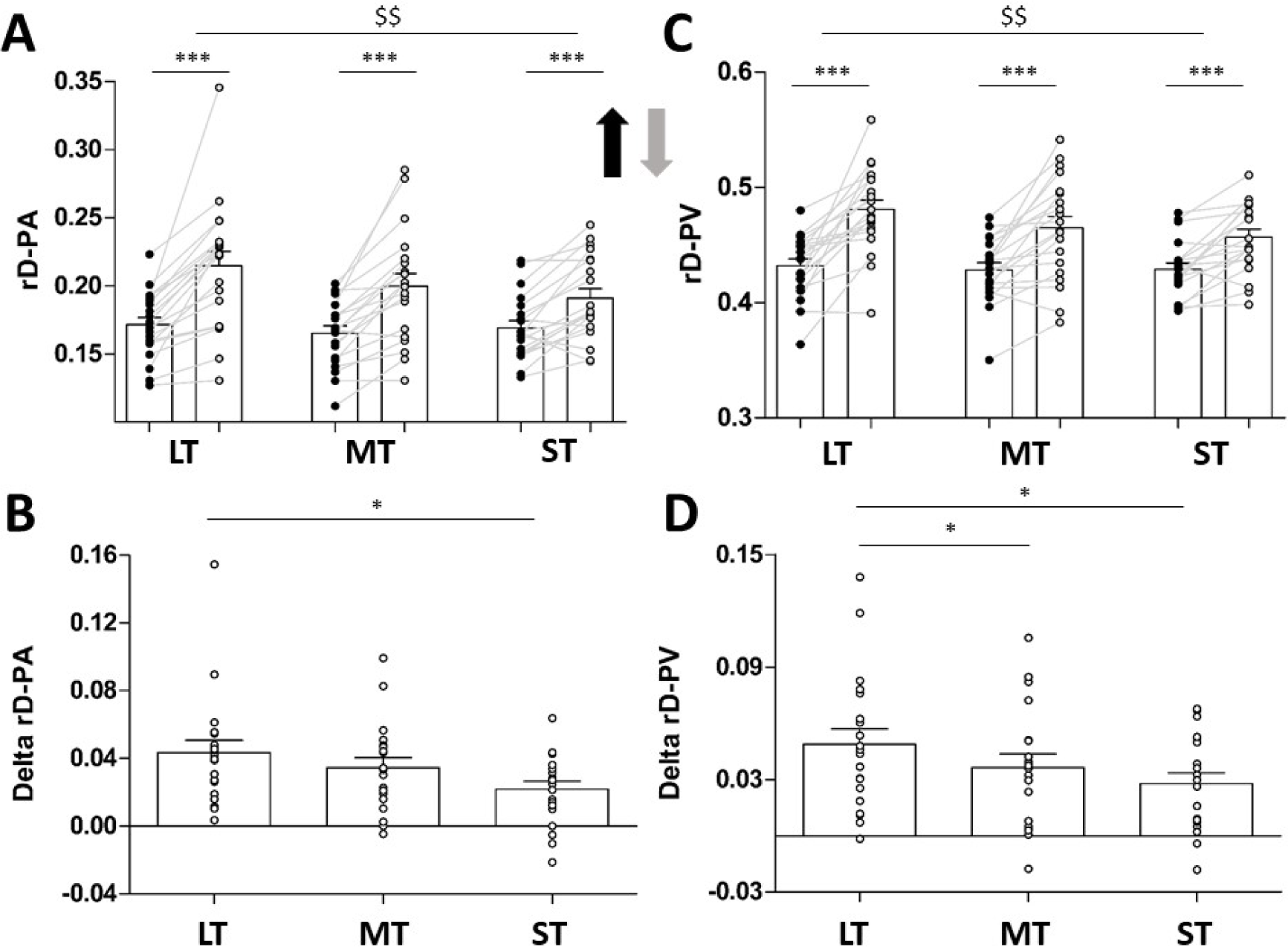
A. Mean (±SD) relative Duration to Peak Acceleration (rD-PA) in upward (black) and downward (grey) movements for all target size conditions. Dots represent individual values of all participants. B. Mean (±SD) values of Delta rD-PA (computed for each participant as: rD-PA_Down_-rD-PA_Up_). Dots represent individual values of all participants. C. Mean (±SD) relative Duration to Peak Velocity (rD-PV) in upward (black) and downward (grey) movements for all target size conditions. Dots represent individual values of all participants. D. Mean (±SD) values of Delta rD-PV (computed for each participant as: rD-PA_Down_-rD-PA_Up_). Dots represent individual values of all participants. $$: Significant main effect of target size; p < 0.01 ***: Upward vs. downward difference on post-hoc test: p < 0.001 *: Significant difference on paired t-test : p < 0.05

Additionally, we observed the *target size x direction* interaction effect to be significant (F = 5.21; p = 1.00e-2; η_p_^2^ = 0.22), demonstrating that directional asymmetry (i.e., difference between upward and downward movements) were reduced as target size decreased. To highlight this effect, we computed a directional difference index between directions (Delta rD-PA; computed as Down-Up values for each participant) for each target size condition (Fig.4B). Preplanned bilateral paired *t* tests revealed that the directional difference was higher when participants reached to the LT compared to the ST (t = 2.60; p = 1.75e-2; Cohen’s d = 0.58). We did not observe statistical differences in other comparisons (ST vs. MT: t = 2.03; p = 5.68e-1; Cohen’s d = 0.45; MT vs. LT: t = 1.69; p = 1.07e-1; Cohen’s d = 0.38).

##### rD-PV

Mean (± SD) rD-PV in each condition is displayed in Fig. 4C. rD-PV was also affected by both *target size* (F = 5.69; p = 6.88e-3; η_p_^2^ = 0.23) and *direction* (F = 40.11; p = 4.46e-6; η_p_^2^ = 0.68) effects. *Post-hoc* comparisons revealed that rD-PV was inferior in upward compared to downward movements in all conditions (Fig. 4C; p < 1.44e-4; Cohen’s d > 0.85 for all comparisons). Additionally, rD-PV in LT condition was found to differ from other conditions in downward (LT vs. MT: p = 1.58e-2; Cohen’s d = 0.49; LT vs.ST: p = 2.31e-4; Cohen’s d = 0.74) but not in upward movements (p > 9.71e-1; Cohen’s d < 0.11 for all comparisons). In the same way as rD-PA, directional asymmetries were reduced as target size decreased, as testified by the significant *target size x direction* interaction effect (F = 5.21; p = 9.97e-3; η_p_^2^ = 0.22). Once again, we computed the directional difference (Down-Up; i.e Delta rD-PV) reflecting directional asymmetry for each condition (Fig. 4D). Preplanned bilateral paired *t* tests highlighted that the Delta rD-PV was larger in the LT compared to the ST (t = 2.66; p = 1.54e-2; Cohen’s d = 0.60) and the MT (t = 2.75; p = 1.27e-2; Cohen’s d = 0.62). No statistical difference was observed between ST and MT conditions (t = 1.26; p = 2.22e-1; Cohen’s d = 0.28).

#### EMG

A qualitative illustration of phasic EMG profiles in all conditions is displayed in Fig. 2C. Phasic EMG patterns during vertical arm movements are characterized by the presence of negative epochs during the deceleration phase of upward movements and during the acceleration phase of downward movements (Gaveau et al., 2021; Poirier et al., 2022, 2023b, 2023a). This negativity reflects the fact that the current EMG activity is inferior than the activity necessary to compensate gravity torque, thereby demonstrating the harnessing of gravity’s effects by the CNS. These negative epochs are now thought to represent a marker of the gravity-related motor optimization (Gaveau et al., 2021; Poirier et al., 2022, 2023b, 2023a).

The negativity has shown to be present in all movement speeds for the downward direction but only in fast movements in the upward direction (Poirier et al., 2023a). Here, because participants were unconstrained concerning movement velocity, we detected negative epochs in all participants and 92.7% of trials overall during downward movements but in 9/20 participants and 14.5% of trials only during upward movements. This is due to the fact that some participants moved too slow to produce negativity.

In this section we analyze 2 parameters: i) negativity occurrence, reflecting the proportion of trials in which negativity was detected in a condition and ii) negativity index, reflecting the magnitude of the negativity in each condition.

##### Negativity Occurrence

Fig. 5A displays mean (± SD) negativity occurrence for each condition. As expected, negativity occurrence was strongly influenced by *direction* (F = 197.11; p = 1.75e-11; η_p_^2^ = 0.91). All participants exhibited higher negativity occurrence values for downward compared to upward movements. We also observed a *target size* effect (F = 3.96; p = 2.75e-2; η_p_^2^ = 0.17): negativity occurrence was reduced as target size decreased (Fig. 5A). *Target size x direction* (F = 0.33; p = 7.20e-1; η_p_^2^ = 0.02) interaction effect was not found to be significant.

**Figure.5.**
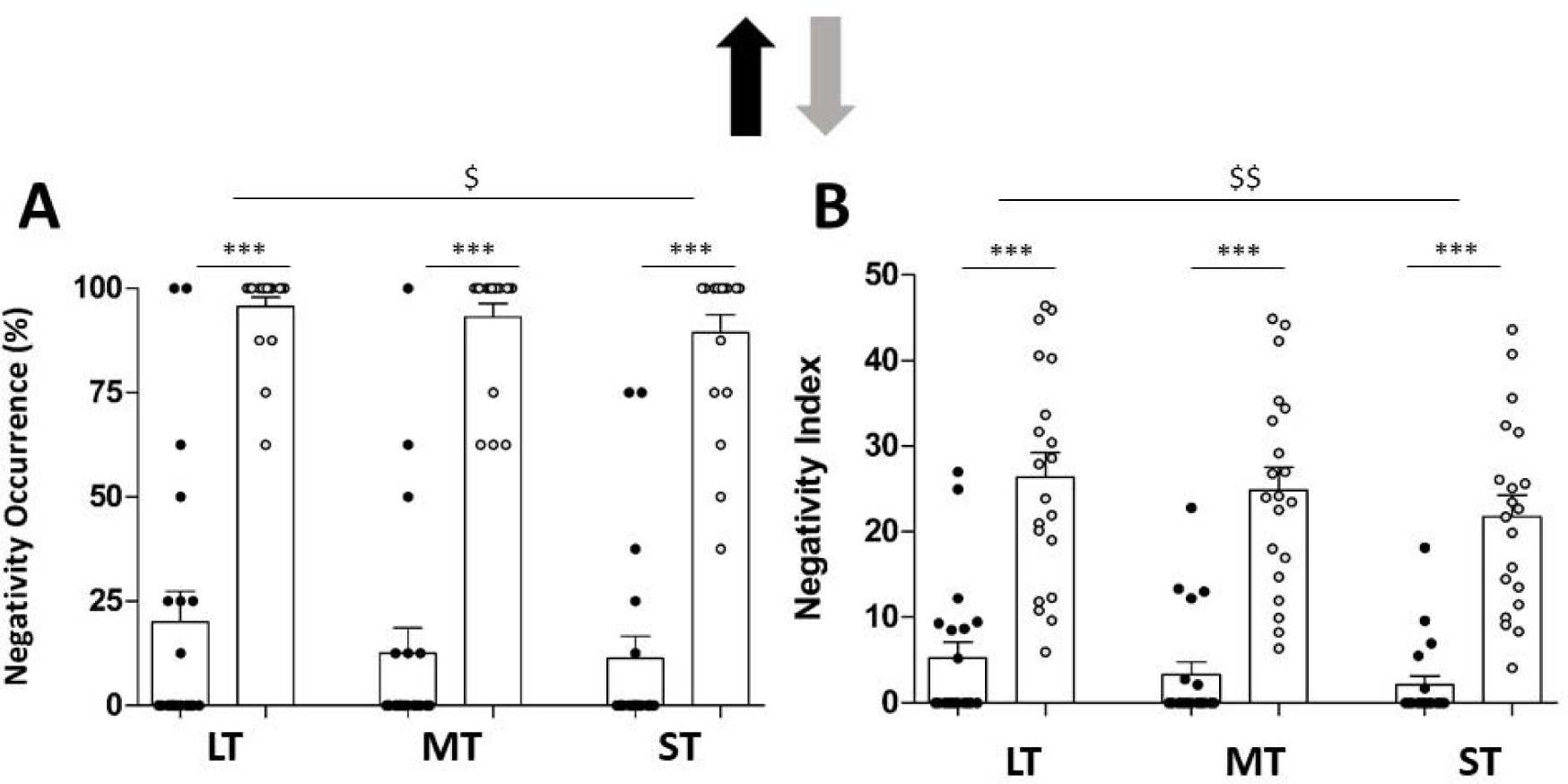
A. Mean (+SD) negativity occurrence for upward and downward movements across target size conditions. Dots represent individual values. B. Mean (+SD) negativity index for upward and downward movements across target size conditions. Dots represent individual values. $: Significant main effect of target size; p < 0.05 $$: Significant main effect of target size; p < 0.01 ***: upward vs. downward difference on post-hoc test: p < 0.001

##### Negativity Index

Fig. 5B displays mean (± SD) negativity index for each condition. Our analysis showed that both *direction* (F = 89.62; p = 1.26e-8; η_p_^2^ = 0.83) and *target size* (F = 7.83; p = 1.42e-3; η_p_^2^ = 0.29) significantly impacted negativity index. However, *target size x direction* interaction effect did not reach significance (F = 1.11; p = 3.40e-1; η_p_^2^ = 0.06). Post-hoc comparisons revealed that negativity index was inferior for upward compared to downward movements in all target size conditions (p < 1.40e-4; Cohen’s d > 2.04 in each case; Fig. 5B). Furthermore, negativity index for downward movements was smaller in the ST condition compared to the MT and LT conditions (p = 2.15e-2; Cohen’s d = 0.32 and p = 3.57e-4; Cohen’s d = 0.47, respectively), but did not differ between MT and LT conditions (p = 6.20e-1; Cohen’s d = 0.15). In upward movements, negativity index was significantly reduced for the ST compared to the LT condition (p = 1.77e-2; Cohen’s d = 0.33), but other comparisons were not significant (p > 3.05e-1; Cohen’s d < 0.21 for both comparisons).

### Relationships between movement duration, effort markers, and variable error

Our results show that the markers of gravity-related effort optimization (i.e., directional asymmetries and negativity phases) were reduced when participants reached to small target sizes compared to large target size. This finding is consistent with a downweighing of the effort cost in favour of the kinematic cost to reduce end-point variability in the case of target downsizing. To confirm this hypothesis, we analysed the relationship between total variability and both markers of gravity-related motor planning among conditions using repeated measures correlations (Bakdash and Marusich, 2017). Because movement duration and coactivation were previously shown to influence end-point variability (Fitts, 1954; Gribble et al., 2003; Missenard and Fernandez, 2011), we also investigated their relationship with variable error.

The relationship between movement duration and total variability was not found to be significant (r_rm_(39) = −0.25, 95% CI [-0.521, 0.057], p = 1.08e-1; Fig. 6A). However, the coactivation index was negatively correlated to total variability (r_rm_(39) = −0.35, 95% CI [-0.597, −0.052], p = 2.30e-2; Fig.6B), highlighting that reduced end-point variability was associated with increased coactivation. Additionally, both Delta rD-PV (r_rm_(39) = 0.31, 95% CI [0.004, 0.565], p = 4.80e-2; Fig.6C) and negativity index (r_rm_(39) = 0.53, 95% CI [0.268, 0.721], p = 3.49e-4; Fig6D) were positively associated with total variability: participants exhibited reduced directional asymmetries and negativity during conditions in which end-point variability was smaller, supporting the hypothesis of a reweighing of effort and kinematic costs.

**Figure.6.**
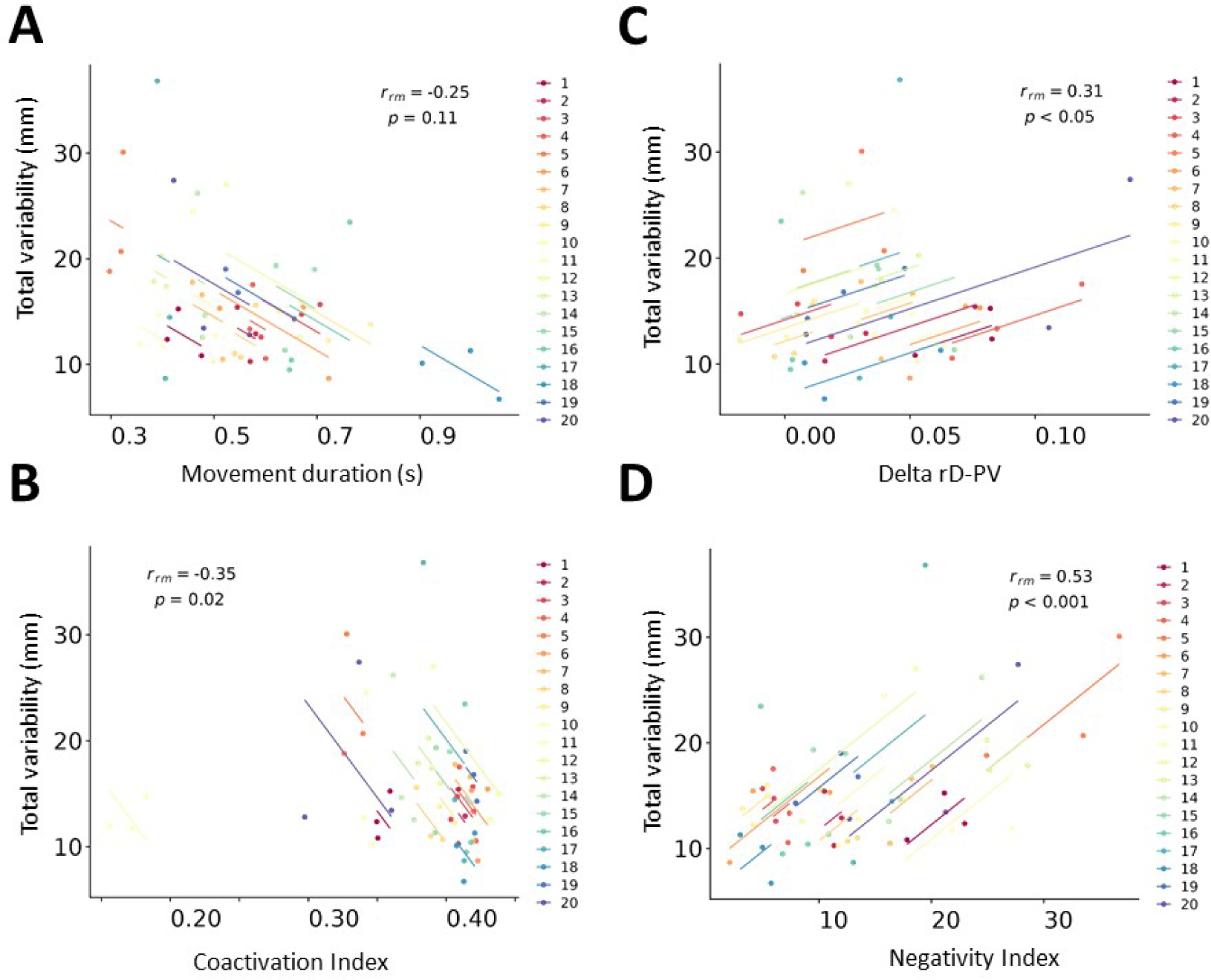
Repeated measures correlation between total variability and movement duration (A), coactivation index (B), delta rD-PV (C), and negativity index (D). Each dot represents a condition and each color represents a participant.

## Discussion

In the present study, we probed the effect of varying accuracy constraints on the gravity-related optimization process. To this aim, we compared the motor signatures of this process (i.e., Directional asymmetries and phasic EMG negativity) in vertical arm pointing movement to a large (LT), a medium (MT) and a small (ST) target. We found a reorganization of motor patterns between the different target size conditions. Our main finding was that directional differences on velocity profiles and negativity index (in downward movements) decreased when target size was reduced. Overall, these results demonstrate that gravity-related effort optimization process is affected by accuracy constraints. More generally, these observations can be interpreted as a reweighting of effort and kinematic costs accordingly to task demands.

First of all, our results concerning the kinematic and EMG patterns of single degree of freedom vertical arm movements are consistent with previous investigations supporting the gravity-related effort-optimization hypothesis (Papaxanthis et al., 2005; Gentili et al., 2007; Le Seac’h and McIntyre, 2007; Gaveau et al., 2014, 2016; Yamamoto and Kushiro, 2014; Gaveau et al., 2021; Yamamoto et al., 2016, 2019; Hondzinski et al., 2016; Poirier et al., 2023a, 2020, 2022, 2023b). We found that relative durations to both peak acceleration and peak velocity were observed shorter for upward compared to downward movements. Additionally, we found direction specific negative phases on the antigravity muscle’s phasic EMG profiles. Precisely, these epochs were found on the acceleration phase of downward movements and the deceleration phase of upward movements. A notable difference is the smaller proportion of trials in which negativity was detected for upward movements: 12.5% in the present study whereas it was around 70-80% in previous researches (Gaveau et al., 2021; Poirier et al., 2022). This is due to the fact that these studies investigated fast movements while movement speed was not constrained in the present study. We recently demonstrated that the generation of substantial inertial torque in the acceleration phase is a necessary condition to the presence of negativity in the deceleration phase of upward movements, because it would otherwise compromise the movement’s integrity (Poirier et al., 2023a). Thus, upward movements must be performed at a certain minimum speed for the CNS to exploit gravity’s effects (around 500ms for the amplitude set here). As the movement velocity was not constrained here, many participants moved too slow to exhibit negativity.

We investigated the effects of varying the target size on motor strategies. As accuracy demands increased, we observed the well-known reorganization of motor patterns: higher movement duration, reduced end-point variability and increased muscular coactivation (Fitts, 1954; Soechting, 1984; Gribble et al., 2003; Osu et al., 2004; Missenard and Fernandez, 2011).

Specifically, we took advantage of previous knowledge to probe the effect of accuracy constraints of gravity-related motor optimization process. We found that, for both rD-PA and rD-PV, the size of directional asymmetries was reduced as target size decreased. Additionally, the negativity index (quantifying the amount of negativity in each condition) was also shown to be decreased when participants reached to smaller targets. These results clearly demonstrate that the gravity-related effort-optimization process is affected by accuracy demands. Optimal control simulations have explained these motor signatures as the result of the minimization of a composite cost function combining effort and jerk (Berret et al., 2008; Gaveau et al., 2014, 2016, 2021). According to these simulations, the size of these markers is proportional to the relative weight of effort in the composite cost (Gaveau et al., 2016). Therefore, our results can be interpreted as a downweighing of the effort-related term in the composite cost. From a behavioural perspective, this signifies that the CNS places a lesser emphasis on effort minimization when accuracy demands are high. The greater coactivation that we observed for smaller targets corroborates this interpretation, since this strategy is known to be effort-consuming (Latash, 2018). In this way, our results support previous studies suggesting that the CNS is able to appropriately modulate the relative importance of motor costs when facing varying task demands (Osu et al., 2004; Liu and Todorov, 2007).

An important question regarding this interpretation concerns the effect of movement duration on these results. Here, we found that participants moved slightly slower as target size was reduced. As stated above, we previously demonstrated that negativity was strongly influenced by movement duration, with faster movements exhibiting increased negativity compared to slower movements (Poirier et al., 2023a). Thus, an alternative interpretation could be that the decrease in size of gravity-related effort optimization markers is not due to target size reduction *per se*, but to the slight increase of movement duration following target size reduction. We invoke two elements to rule out this hypothesis. First, the difference in movement duration between condition, although significant, is much smaller than in our previous study (in average 50ms between the fastest and the slowest conditions here). Therefore, the influence of movement duration on gravity-related effort optimization markers, if any, should be minimal. Secondly, directional asymmetries do not follow the same relationship with movement duration than negativity. Previous researches found either no speed effect on directional asymmetries (Gaveau and Papaxanthis, 2011) or increased directional asymmetries for slow compared to fast movements (Poirier et al., 2020). Thus, if differences in movement duration were responsible for the observed differences in directional asymmetries between conditions, we should have observed higher directional asymmetries for smaller targets, which is not the case.

Our results also add to the knowledge concerning the relationship between movement speed, effort, and accuracy. Previous investigations have identified two control strategies regarding increasing accuracy demands: reducing movement time or increasing coactivation (Fitts, 1954; Gribble et al., 2003; Osu et al., 2004; Missenard and Fernandez, 2011). The increase in muscle coactivation was found to be the strategy individuals used to cope with higher accuracy requirement when movement time was constrained (Gribble et al., 2003; Osu et al., 2004). Inversely in the case of unconstrained movement time (i.e., move as fast and accurately as possible), participants were observed not to increase coactivation levels but chose to move slower (Missenard and Fernandez, 2011). Here, as target size decreased, participants moved slightly slower but also increased coactivation and decreased the size of gravity-related effort optimization signatures. Repeated measures correlations showed that the reduction in end-point variability was associated with: i) a reduction of directional asymmetries and negativity and ii) an increase of coactivation level, but was not associated with movement time. As participants were totally free concerning movement time (they were asked to move at their preferential speed), it may seem surprising that they used such an effort-consuming strategy to reduce end-point variability instead of simply moving slower. A possible explanation lies in the cost of time theory (Shadmehr et al., 2010; Berret and Jean, 2016). This theory stems from the proposition that the purpose of any action would be to put the neural system in a more rewarding state (Shadmehr et al., 2010). Because neuroeconomical studies demonstrated the CNS tendency to discount the value of future reward, slower movements may be undesirable because the passage of time entails a cost per se. Thus, movement duration can be considered as the outcome of the minimization of a subjective weighting between a cost of time and a cost of movement, depending of expected reward. In tasks where reward is not explicit (as in our task), multiple studies showed that movement duration can be appropriately captured by a common tradeoff between time and effort (Shadmehr et al., 2010; Berret and Jean, 2016; Berret and Baud-Bovy, 2022; Carlisle et al., 2023; Verdel et al., 2023). The existence of this cost of time can explain why our participants did not choose to move slower and adopted a more effort-consuming strategy. The reorganization of motor patterns that we observed when accuracy demands increased can in this way be interpreted as an optimal tradeoff between time and effort.

## References

Bakdash, J. Z., and Marusich, L. R. (2017). Repeated Measures Correlation. Front. Psychol. 8, 456. doi:10.3389/fpsyg.2017.00456.

Berret, B., and Baud-Bovy, G. (2022). Evidence for a cost of time in the invigoration of isometric reaching movements. J. Neurophysiol. 127, 689–701. doi:10.1152/JN.00536.2021/ASSET/IMAGES/LARGE/JN.00536.2021_F009.JPEG.

Berret, B., Chiovetto, E., Nori, F., and Pozzo, T. (2011). Evidence for composite cost functions in arm movement planning: An inverse optimal control approach. PLoS Comput. Biol. 7, e1002183. doi:10.1371/journal.pcbi.1002183.

Berret, B., Darlot, C., Jean, F., Pozzo, T., Papaxanthis, C., and Gauthier, J. P. (2008). The inactivation principle: Mathematical solutions minimizing the absolute work and biological implications for the planning of arm movements. PLoS Comput. Biol. 4, e1000194. doi:10.1371/journal.pcbi.1000194.

Berret, B., and Jean, F. (2016). Why Don’t We Move Slower? The Value of Time in the Neural Control of Action. J. Neurosci. 36, 1056–1070. doi:10.1523/JNEUROSCI.1921-15.2016.

Buneo, C. A., Soechting, J. F., and Flanders, M. (1994). Muscle activation patterns for reaching: The representation of distance and time. J. Neurophysiol. 71, 1546–1558. doi:10.1152/jn.1994.71.4.1546.

Carlisle, R., Elife, A. K.-, and 2023, undefined (2023). Optimization of energy and time predicts dynamic speeds for human walking. elifesciences.org 12, 81939. doi:10.7554/eLife.

D’Avella, A., Fernandez, L., Portone, A., and Lacquaniti, F. (2008). Modulation of phasic and tonic muscle synergies with reaching direction and speed. J. Neurophysiol. 100, 1433–1454. doi:10.1152/jn.01377.2007.

D’Avella, A., Portone, A., Fernandez, L., and Lacquaniti, F. (2006). Control of fast-reaching movements by muscle synergy combinations. J. Neurosci. 26, 7791–7810. doi:10.1523/JNEUROSCI.0830-06.2006.

Fitts, P. M. (1954). The information capacity of the human motor system in controlling the amplitude of movement. J. Exp. Psychol. 47, 381–391. doi:10.1037/H0055392.

Flanders, M., and Herrmann, U. (1992). Two components of muscle activation: Scaling with the speed of arm movement. J. Neurophysiol. 67, 931–943. doi:10.1152/jn.1992.67.4.931.

Flanders, M., Pellegrini, J. J., and Soechting, J. F. (1994). Spatial/temporal characteristics of a motor pattern for reaching. J. Neurophysiol. 71, 811–813. doi:10.1152/jn.1994.71.2.811.

Franklin, D. W., and Wolpert, D. M. (2011). Computational mechanisms of sensorimotor control. Neuron 72, 425–442. doi:10.1016/j.neuron.2011.10.006.

Gaveau, J., Berret, B., Angelaki, D. E., and Papaxanthis, C. (2016). Direction-dependent arm kinematics reveal optimal integration of gravity cues. Elife 5. doi:10.7554/eLife.16394.

Gaveau, J., Berret, B., Demougeot, L., Fadiga, L., Pozzo, T., and Papaxanthis, C. (2014). Energy-related optimal control accounts for gravitational load: comparing shoulder, elbow, and wrist rotations. J. Neurophysiol. 111, 4–16. doi:10.1152/jn.01029.2012.

Gaveau, J., Grospretre, S., Berret, B., Angelaki, D. E., and Papaxanthis, C. (2021). A cross-species neural integration of gravity for motor optimization. Sci. Adv. 7, eabf7800. doi:10.1126/sciadv.abf7800.

Gaveau, J., and Papaxanthis, C. (2011). The temporal structure of vertical arm movements. PLoS One 6, e22045. doi:10.1371/journal.pone.0022045.

Gentili, R., Cahouet, V., and Papaxanthis, C. (2007). Motor planning of arm movements is direction-dependent in the gravity field. Neuroscience 145, 20–32. doi:10.1016/j.neuroscience.2006.11.035.

Gielen, S. (2009). Review of models for the generation of multi-joint movements in 3-D. Adv. Exp. Med. Biol. 629, 523–550. doi:10.1007/978-0-387-77064-2_28.

Gribble, P. L., Mullin, L. I., Cothros, N., and Mattar, A. (2003). Role of cocontraction in arm movement accuracy. J. Neurophysiol. 89, 2396–2405. doi:10.1152/jn.01020.2002.

Healy, C. M., Berniker, M., and Ahmed, A. A. (2023). Learning vs. minding: How subjective costs can mask motor learning. PLoS One 18, e0282693. doi:10.1371/JOURNAL.PONE.0282693.

Hondzinski, J. M., Soebbing, C. M., French, A. E., and Winges, S. A. (2016). Different damping responses explain vertical endpoint error differences between visual conditions. Exp. Brain Res. 234, 1575–1587. doi:10.1007/s00221-015-4546-8.

Jordan, M. I., and Wolpert, D. M. (1999). Computational motor control. Cogn. Neurosci. 601, 597–609. doi:10.1073/pnas.0914906108.

Latash, M. L. (2018). Muscle coactivation: Definitions, mechanisms, and functions. J. Neurophysiol. 120, 88–104. doi:10.1152/jn.00084.2018.

Le Seac’h, A. B., and McIntyre, J. (2007). Multimodal reference frame for the planning of vertical arms movements. Neurosci. Lett. 423, 211–215. doi:10.1016/j.neulet.2007.07.034.

Liu, D., and Todorov, E. (2007). Evidence for the Flexible Sensorimotor Strategies Predicted by Optimal Feedback Control. J. Neurosci. 27, 9354–9368. doi:10.1523/JNEUROSCI.1110-06.2007.

Missenard, O., and Fernandez, L. (2011). Moving faster while preserving accuracy. Neuroscience 197, 233–241. doi:10.1016/J.NEUROSCIENCE.2011.09.020.

Missenard, O., Mottet, D., and Perrey, S. (2008). The role of cocontraction in the impairment of movement accuracy with fatigue. Exp. Brain Res. 185, 151–156. doi:10.1007/S00221-007-1264-X/FIGURES/3.

Mombaur, K., Truong, A., and Laumond, J. P. (2010). From human to humanoid locomotion-an inverse optimal control approach. Auton. Robots 28, 369–383. doi:10.1007/S10514-009-9170-7/METRICS.

Oldfield, R. C. (1971). The assessment and analysis of handedness: The Edinburgh inventory. Neuropsychologia 9, 97–113. doi:10.1016/0028-3932(71)90067-4.

Osu, R., Kamimura, N., Iwasaki, H., Nakano, E., Harris, C. M., Wada, Y., et al. (2004). Optimal impedance control for task achievement in the presence of signal-dependent noise. J. Neurophysiol. 92, 1199–1215. doi:10.1152/JN.00519.2003/ASSET/IMAGES/LARGE/Z9K0080439950011.JPEG.

Papaxanthis, C., Pozzo, T., and McIntyre, J. (2005). Kinematic and dynamic processes for the control of pointing movements in humans revealed by short-term exposure to microgravity. Neuroscience 135, 371–383. doi:10.1016/j.neuroscience.2005.06.063.

Poirier, G., Mourey, F., Sirandre, C., Papaxanthis, C., and Gaveau, J. (2023a). Speed-dependent optimization of gravity effects for motor control. bioRxiv, 2023.03.14.532654. doi:10.1101/2023.03.14.532654.

Poirier, G., Papaxanthis, C., Lebigre, M., Juranville, A., Mathieu, R., Savoye-Laurens, T., et al. (2023b). Aging decreases the lateralization of gravity-related effort minimization during vertical arm movements. bioRxiv, 2021.10.26.465988. doi:10.1101/2021.10.26.465988.

Poirier, G., Papaxanthis, C., Mourey, F., and Gaveau, J. (2020). Motor Planning of Vertical Arm Movements in Healthy Older Adults: Does Effort Minimization Persist With Aging? Front. Aging Neurosci. 12. doi:10.3389/fnagi.2020.00037.

Poirier, G., Papaxanthis, C., Mourey, F., Lebigre, M., and Gaveau, J. (2022). Muscle effort is best minimized by the right-dominant arm in the gravity field. J. Neurophysiol. 127, 1117–1126. doi:10.1152/jn.00324.2021.

Schmidt, R., and Lee, T. (2019). Motor Control and Learning: A Behavioral Emphasis, 6th Edition. Human Kinetics doi:10.1249/01.mss.0000550736.00894.1f.

Scott, S. H. (2004). Optimal feedback control and the neural basis of volitional motor control. Nat. Rev. Neurosci. 5, 532–544. doi:10.1038/nrn1427.

Shadmehr, R., De Xivry, J. J. O., Xu-Wilson, M., and Shih, T. Y. (2010). Temporal Discounting of Reward and the Cost of Time in Motor Control. J. Neurosci. 30, 10507–10516. doi:10.1523/JNEUROSCI.1343-10.2010.

Soechting, J. F. (1984). Effect of target size on spatial and temporal characteristics of a pointing movement in man. Exp. Brain Res. 54, 121–132. doi:10.1007/BF00235824/METRICS.

Tanis, D., Calalo, J. A., Cashaback, J. G. A., and Kurtzer, I. L. (2023). Accuracy and effort costs together lead to temporal asynchrony of multiple motor commands. J. Neurophysiol. 129, 1–6. doi:10.1152/JN.00435.2022/ASSET/IMAGES/LARGE/JN.00435.2022_F003.JPEG.

Todorov, E. (2004). Optimality principles in sensorimotor control. Nat. Neurosci. 7, 907–915. doi:10.1038/nn1309.

Verdel, D., Bruneau, O., Sahm, G., Vignais, N., and Berret, B. (2023). The value of time in the invigoration of human movements when interacting with a robotic exoskeleton. Sci. Adv. 9. doi:10.1126/SCIADV.ADH9533.

Vu, V. H., Isableu, B., and Berret, B. (2016). On the nature of motor planning variables during arm pointing movement: Compositeness and speed dependence. Neuroscience 328, 127–146. doi:10.1016/j.neuroscience.2016.04.027.

Wolpert, D. M. (1997). Computational approaches to motor control. Trends Cogn. Sci. 1, 209–216. doi:10.1016/S1364-6613(97)01070-X.

Yamamoto, S., Fujii, K., Zippo, K., Kushiro, K., and Araki, M. (2019). The kinetic mechanisms of vertical pointing movements. Heliyon 5, e02012. doi:10.1016/J.HELIYON.2019.E02012.

Yamamoto, S., and Kushiro, K. (2014). Direction-dependent differences in temporal kinematics for vertical prehension movements. Exp. Brain Res. 232, 703–711. doi:10.1007/s00221-013-3783-y.

Yamamoto, S., Shiraki, Y., Uehara, S., and Kushiro, K. (2016). Motor control of downward object-transport movements with precision grip by object weight. Somatosens. Mot. Res. 33, 130–136. doi:10.1080/08990220.2016.1203304.

